# Engineered lysins to modulate human gut microbiome communities

**DOI:** 10.1101/2024.05.14.594189

**Authors:** Juan R. Diaz Rodriguez, Wenbo Lu, James M. Papadopoulos, Ophelia S. Venturelli, Philip A. Romero

## Abstract

The ability to precisely control the human gut microbiome’s composition, metabolic networks, and host interactions would revolutionize medicine and how we treat human disease. Microbiome remodeling approaches such as probiotics, prebiotics, antibiotics, and fecal microbiota transplantation are global perturbations with consequences that are difficult to predict. Lysins are enzymes produced by bacteriophages and bacteria that have exceptional specificity to lyse bacterial cells and could be applied for targeted microbiome remodeling. In this work we demonstrate the ability of lysins to target diverse human gut commensal bacteria, how domain recombination can give rise to chimeric lysins with altered species preferences, and how engineered lysins can directly remodel the structure and function of synthetic human gut microbiomes. Lysins represent a general platform that can be rapidly tailored through protein engineering to control complex microbial ecosystems.

## Introduction

The human gut microbiome is a complex ecosystem that plays a crucial role in human health and disease. The composition and function of the gut microbiome have been associated with metabolic, immunological, and neurological illnesses. Moreover, the gut microbiome has been recognized as a key player in polysaccharide digestion, vitamin and nutrient biosynthesis, colonization resistance, and immune system modulation [1], [2]. Metabolites produced by the microbiome are known to affect host physiology [3]. For example, decreased butyrate production by the microbiome has been linked to inflammatory bowel disease, obesity, and metabolic disorders [4]. Altered community composition and metabolic functions of the gut microbiome in response to certain environmental factors or host dysfunctions can contribute to multiple human diseases [5], [6].

There is growing interest in developing novel precision microbiome engineering strategies that alter key functions of the gut microbiome to our benefit [7]. Microbiome remodeling has previously been achieved using approaches such as probiotics, prebiotics, and fecal microbiota transplantation. However, these methods have limitations regarding specificity, efficacy, and long-term stability [8]. Recently, direct remodeling has been proposed as a more targeted approach by using antimicrobial peptides (AMPs) and has successfully ameliorated the development of atherosclerosis in a mouse model [9]. This approach, however, faces challenges due to AMPs’ broad spectrum resulting in off-target global microbiome perturbations. Therefore, new control knobs for shaping the composition and function of gut microbiota are needed.

Lysins are enzymes produced by bacteriophages and bacteria that specifically target and lyse bacterial cells and offer an alternate approach to microbiome remodeling. Gram-positive lysins are hydrolytic enzymes that cleave the peptidoglycan cell wall, resulting in lysis and death. These proteins are modular, with specificity conferred by separate enzymatic and cell wall binding domains and have previously been engineered with altered specificity and enhanced activity against target bacteria [10]. Furthermore, they can be modified to target gram-negative bacteria by adding outer membrane permeating peptides [11], [12]. The potential for engineering specificity and flexibility in lysins makes them valuable tools for modulating gut microbiome composition.

In this study, we explored the ability of lysins to precisely control the structure and function of the human gut microbiome. We designed a panel of chimeric lysins by domain shuffling and screened the chimeric lysins for their ability to inhibit the growth of human commensal bacteria. The chimeric lysins showed varied specificity across diverse and prevalent human gut bacteria. We identified sequence patterns and domains that drive activity and specificity toward individual species. We next applied our engineered lysins to synthetic gut microbiome communities and measured the system’s biological responses. We found lysins are potent modulators of the community structure and caused alterations in key fermentation end products associated with human health. This work establishes lysins as a general platform for controlling the human gut microbiome.

## Results

### Chimeric lysins show varied specificity across human gut commensals

Lysin specificity is dictated by both enzyme activity domains (EADs) and cell-wall binding domains (CBDs). We wanted to explore if recombining diverse EAD-CBD combinations would give rise to new lysins with novel specificity profiles. We started with Enterolysin A (EnlA) from *E. faecalis* because it is known to have broad activity across intestinal flora from enterococci, pediococci, lactococci, and lactobacilli species.

We searched the UniRef100 sequence database for diverse sequences homologous to EnlA’s EAD and CBD domains (**Fig. 1a**). We identified two EAD domains from lysins belonging to *Lactococcus lactis* and *Enterococcus avium*, which we refer to as EAD2 and EAD3, and that both share 56% sequence identity to EnlA. Like EnlA’s EAD, these new domains are annotated as M23 metallopeptidase family peptidoglycan hydrolases. We also searched UniRef100 for sequences related to EnlA’s CBD domain and identified two CBD domains from Lactococcus phage *asccphi28* and *Lactococcus lactis* (CBD2 and CBD3) that share 60% and 61% sequence identity to EnlA, respectively. CBD domains are highly variable and not well annotated, and our attempts to classify the CBDs through InterPro found no associated protein families or motifs. Nevertheless, both CBD2 and CBD3 displayed adequate similarity to the query EnlA CBD in terms of identity and coverage, indicating their potential for creating functional chimeric lysins with diverse sequences.

**Figure 1.**
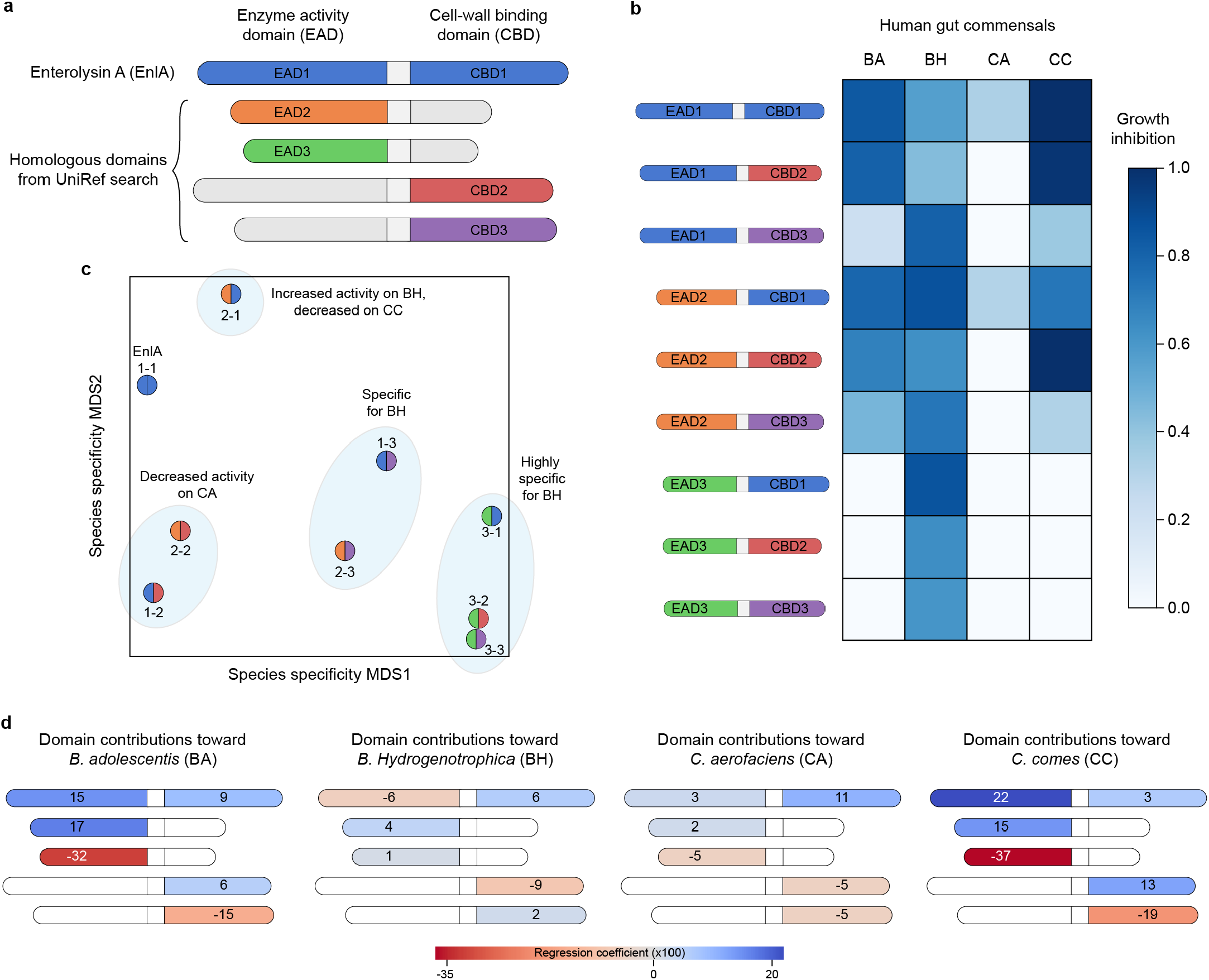
Design and screening of chimeric lysins. (**a**) Lysins consist of enzyme activity domains (EADs) and cell-wall binding domains (CBDs). We identified a diverse set of three EADs and three CBDs that can be shuffled to generate nice chimeric lysins. (**b**) We screened all nine chimeric lysins against a panel of four human gut commensal species. Growth inhibition was calculated by comparing the calculating the area under the growth curves (AUCs) for lysin and no lysin samples. (**c**) Visualization of each chimeric lysins specificity profile from panel b. We used multidimensional scaling (MDS) to reduce the 4-dimensional space to 2D for visualization. Each lysin is colored according to its constituent domains. The lysins cluster into distinct groups based on their specificity. (**d**) We used linear regression to estimate how each domain contributes to activity on each species. The domains are organized according to the layout in panel a. Blue colors indicate increased activity, while red colors indicate reduced activity.

We cloned all nine combinations of domains (3 EADs x 3 CBDs) and expressed the chimeric lysins in *E. coli* (**Supplementary Table 1**). We screened the chimeric lysins for growth inhibition activity across a diverse panel of human gut commensals including *Bifidobacterium adolescentis* (BA), *Blautia Hydrogenotrophica* (BH), *Collinsella aerofaciens* (CA), and *Coprococcus comes* (CC) (**Fig. 1b**). The chimeric lysins displayed markedly different species specificity from wildtype EnlA and each other, suggesting the individual domains and their interactions contribute to microbe recognition and lysis. We visualized the specificity profiles of the nine lysins using multidimensional scaling (MDS) and found they clustered based on activity and specificity (**Fig. 1c**). The most prominent effect came from EAD3, which substantially decreased activity on BA, CA, and CC, resulting in lysins highly specific for BH. The chimera EAD2-CBD1 was interesting because it showed activity on BA, BH, CA, but decreased activity on CC relative to EnlA. CC is an important butyrate producing bacteria with known benifical effects on human health [13]. The chimera EAD2-CBD2 was also interesting because it had similar overall activity and specificity to EnlA but was completely inactive on CA, a microbe associated with reduced risk of colon cancer [14]

We used linear regression analysis to estimate how each domain contributes to activity on each of the four human gut commensal species (**Fig. 1d, Supplementary Fig. 1**). Chimeras with EAD1 and EAD2 were active on BA, but those with EAD3 showed no activity. Similarly, CBD1 and CBD2 contribute to activity on BA, while CBD3 decreases activity. All tested chimeric lysins were active on BH and the strongest effects driving activity seem to arise from the CBD domains. For CA, almost all lysins were inactive except those with CBD1. CC looked very similar to BA with both EADs and CBDs contributing to activity. Based on these results, EAD3 seems highly specific for cleaving BH peptidoglycan. Additionally, activity on BH and CA is largely determined by CBDs, while activity on BA and CC is determined by both EADs and CBDs.

### Quantitative assessment of lysin activity against individual members of a synthetic community

The screening experiments in the previous section were performed with *E. coli* lysates and thus did not control for the absolute lysin concentration. We wanted to obtain more quantitative activity measurements to understand how lysin concentration affects growth inhibition across microbial species. We focused on EnlA (EAD1-CBD1) and the chimera Ch2-2 (EAD2-CBD2) due to their high expression and broad activity. We evaluated these two lysins on a diverse mix of fast growing, slow growing, and butyrate-producing microbes that were chosen in a previous study [15]. These microbes include *Bifidobacterium adolescentis* (BA), *Coprococcus comes* (CC), *Anaerostipes caccae* (AC), *Eubacterium rectale* (ER), and *Roseburia intestinalis* (RI).

We titrated purified lysins against single strains and calculated each microbe’s relative growth at varying lysin concentrations (**Fig. 2**). EnlA showed strong growth inhibition against BA and CC, with maximum activity at 12.5 nM and 100 nM, respectively. EnlA showed moderate growth inhibition against AC with a maximum activity at 100 nM. Ch2-2 showed growth inhibition against BA, CC, and AC, but had overall lower activity relative to EnlA. At higher concentrations beyond 100 nM, EnlA shows preference for BA over CC, while Ch2-2 shows the opposite preference for CC over BA. Both EnlA and Ch2-2 had no activity against ER and RI. The quantitative differences and species preferences between EnlA and Ch2-2 will affect how they behave in a microbial community context.

**Figure 2.**
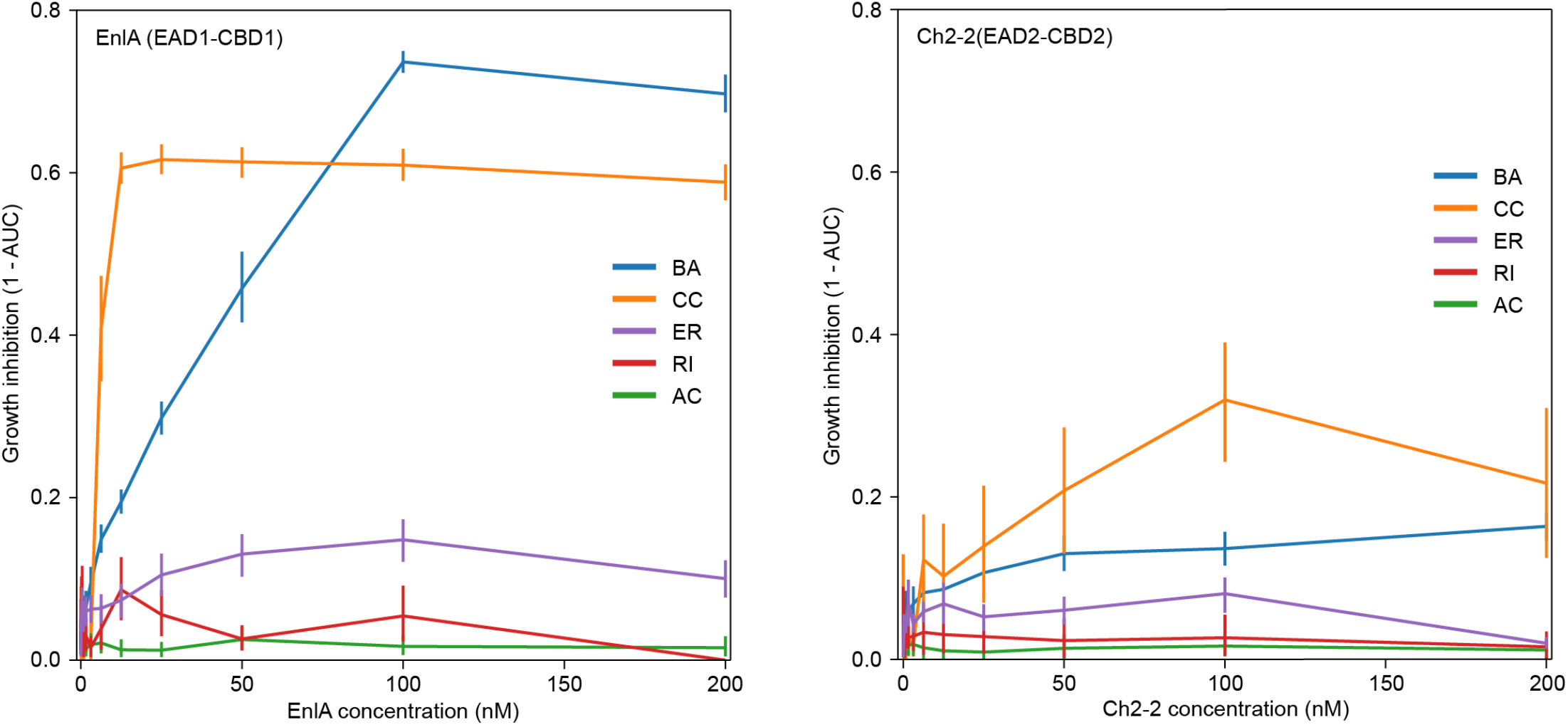
Quantitative titrations of lysins against single species. We titrated purified EnlA (top panel) and Ch2-2 (bottom panel) against monocultures of *Bifidobacterium adolescentis* (BA), *Coprococcus comes* (CC), *Anaerostipes caccae* (AC), *Eubacterium rectale* (ER), and *Roseburia intestinalis* (RI). Growth inhibition was calculated by comparing the calculating the area under the growth curves (AUCs) for lysin and no lysin samples.

### Modulating the structure and function of synthetic gut microbiome communities

Microbial communities are rich ecosystems with many interdependent interactions that shape their behaviors. We wanted to test if our engineered lysins could modulate the structure and function of synthetic gut microbiome communities. We designed two synthetic communities based on a previous study [15] that result in coexistence between species and produce short chain fatty acids and other health-relvant fermentation end products. The first community (COM1) consists of BA and the three butyrate-producers CC, RI and ER. The second community (COM2) consists of BA and the three butyrate-producers CC, RI and AC.

We tested the response of synthetic communities COM1 and COM2 to both lysins EnlA or Ch2-2 and measured abundance of each microbe and the quantity of fermentation end products produced. Increasing lysin concentration generally caused a decrease in overall community density but affected each microbial species differently (**Fig. 3a**). In COM1, there was a marked decrease in the presence of BA and this seems to open a niche for the slow growing RI to expand at high EnlA concentrations. We also found CC was largely unchanged across lysin concentrations despite showing high growth inhibition in single species measurements (**Fig 1 and Fig 2**). In the community context, CC competes with BA and may be less affected by lysins that preferentially lyse BA. 1000 nM EnlA causes complete collapse of COM1, reducing both the community density and diversity. We found COM2 is less affected by lysins, and this seems to be a result of AC having minimal lysin sensitivity. Overall, we see microbes in the community context are less affected by lysins than in single-species experiments. We performed multidimensional scaling (MDS) on the community composition to visualize how lysins reshape the community structure (**Fig. 3b**). We found COM1 and COM2 occupy distinct regions in community space and increasing lysin concentration causes directional changes in the community composition. EnlA and Ch2-2 affect COM1 in a similar direction, but Ch2-2 requires ~10x more protein to achieve a similar effect to EnlA. EnlA and Ch2-2 have more varied effects on COM2, with the lower lysin concentrations moving the communities in different directions. We also measured the fermentation end products produced by the communities (**Fig. 3c**). For COM1, both lysins substantially decreased acetate levels, while increasing both lactate and butyrate. 1000 nM Ch2-2 nearly doubled the buyrate produced by COM1. For COM2, both lysins decreased acetate, increased lactate, and slightly decreased butyrate. These results demonstrate how small quantities of lysins can drastically alter the structure and function of microbial communities.

**Figure 3.**
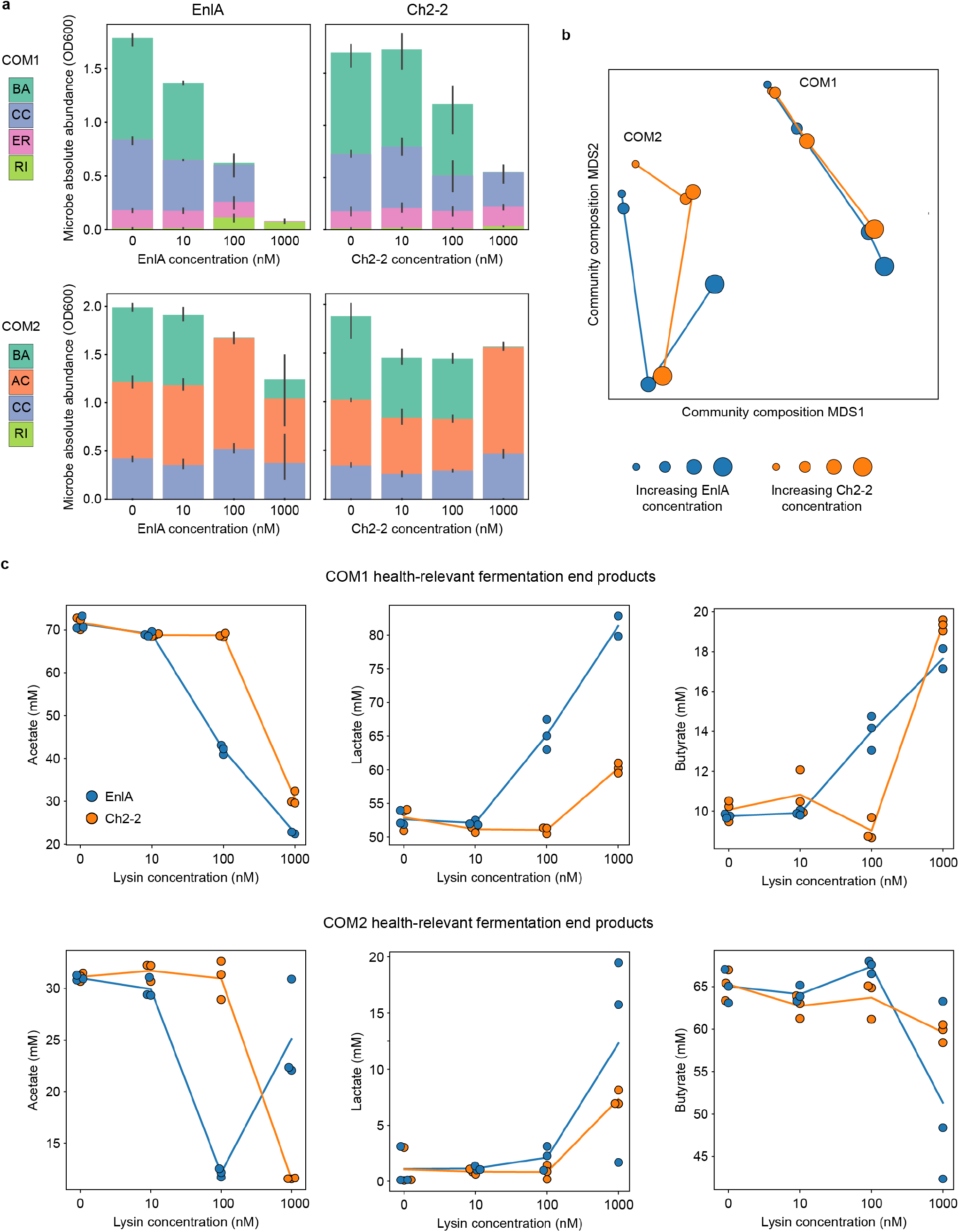
Modulating synthetic gut microbiome communities. (**a**) The community composition of COM1 (top row) and COM2 (bottom row) in response to increasing lysin concentrations. The bars represent absolute abundance of each microbe and the community experiments were performed in triplicate. (**b**) We visualized the response of COM1 and COM2 to increasing lysin concentrations using multidimensional scaling (MDS). The full five-dimensional community (BA, CC, EM, RI, and AC) was reduced to two dimensions. Increasing point size corresponds to increasing lysin concentration. (**c**) Fermentation end products of COM1 (top row) and COM2 (bottom row) at varying lysin concentrations. Each community was analyzed with three biological replicates shown as individual points and the lines correspond to the mean fatty acid concentration.

## Discussion

The human gut microbiome plays essential roles in human health and disease, and strategies to modulate and control this complex ecosystem could be used as therapeutics. Our work demonstrated the ability of lysins to target diverse human gut commensal bacteria, how domain recombination can give rise to chimeric lysins with altered species preferences, and how engineered lysins can directly remodel the structure and function of microbial communities. Lysins provide a general platform that can be rapidly tailored through protein engineering to control complex microbial ecosystems.

Our work is the first example using lysins to remodel the gut microbiome. Other targeted methods gut microbiome manipulation include prebiotics [16], [17], probiotics [18], antimicrobials [9], and phage therapy [19]. These approaches have limited potential to specifically target a subset of microbes and tend to broadly modulate the gut microbiome from a dysbiosis state into a healthy state. Lysins offer two levels of specificity, occurring at the cell-wall binding domain (CBD) and the enzyme activity domain (EAD), and thus provide tunable specify from narrow to broad. One challenge in applying lysins and other antimicrobials to control microbiomes is it’s not clear a priori which microbes should be targeted to achieve a desired microbiome state. Interspecies interactions within a community give rise to complex behaviors that are difficult to predict.

There are many previous examples engineering lysins to target diverse bacterial species and treat bacterial infections. For example, work done by Gerstmans et al. developed a lysin domain shuffling and screening platform capable of creating lysins that target multiple strains of the gram-negative pathogen *A. bumanii* [11]. Other work shuffled lysin domains to target Gardella biofilms for treating and preventing bacterial vaginosis [20]. Lysins can additionally be fused to outer membrane permeabilizing (OMP) peptides to bypass the outer membrane and target gram-negative bacteria [11], [21]. The modular structure and broad species range of lysins makes them ideal scaffolds for selectively targeting diverse microbial species.

The EnlA and Ch2-2 lysins showed potent activity to modulate the species structure of the synthetic human gut communities COM1 and COM2. These changes in species composition greatly influenced the short-chain fatty acid and fermentation end product pools. Overall, the lysins reduced acetate, while increasing lactate and butyrate, all of which could have beneficial effects on the host. Microbially produced acetate is known to activate the parasympathetic nervous system, promoting ghrelin secretion and glucose-stimulated insulin secretion, resulting in hyperlipidemia, fatty liver disease, insulin resistance and obesity [22]. Lactate is known to promote gut microbial diversity and gut health through anti-inflammatory and immunomodulatory effects [23]. Butyrate is a primary energy source for colonocytes, promotes a healthy gut barrier, limits pro-inflammatory cytokines, and inhibits oncogenic pathways [24].

We tested our engineered lysins in synthetic gut communities *in vitro*, but we would ultimately want to deliver lysins to the human gut. Oral delivery of proteins presents challenges with passing through the proteolytic and the acidic environment of the digestive tract. We envision engineering probiotic bacteria that secrete lysins and could be delivered orally. These probiotics could include *E. coli* Nissle 1917, lactobacillus, and bacteroides. Engineered probiotic bacteria could reside in the human gut for short or long timeframes and deliver designed lysins directly to the gastrointestinal tract.

Precisely controlling the structure, function, and behavior of the human gut microbiome will create a revolution in medicine and how we treat human disease. This level of control requires a comprehensive understanding of the interactions within and between the microbiome, the host, and the metabolome, and specific control knobs that alter the microbiome in a predictable manner. Furthermore, the microbiome states associated with human health are largely individual-specific and may require personalized microbiome therapies [19]. Lysins and other antimicrobial peptides/proteins provide a framework for targeted modulation of complex microbial ecosystems *in situ* and can be rapidly engineered to tailor the microbiome toward human health.

## Methods

### Bioinformatics search for lysin EAD and CBD domains

The sequence of wild-type enterolysin A (EnlA) was separated into EAD and CBD at the previously identified TP-rich linker at positions 181-194. These domains were then independently queried using jackhmmer[25] against the Uniref100 database to identify distant homologous domains. Query results were further processed by setting cutoffs of less than 70% pairwise identity and over 80% sequence coverage to the query sequence. This was done to ensure complete domains were selected and that they were sufficiently different from the query sequence and to each other. All domains identified that met the filtering criteria were submitted to InterPro for domain classification. For domains that were successfully classified by InterPro, their sequences were further analyzed to confirm the presence of relevant protein family motifs.

### Lysin cloning and protein expression

Enterolysin A and other homologous domain genes were ordered as Gblocks (IDT). All gene inserts were created by PCR with appropriate GoldenGate adapters to such that all possible domain combinations could be assembled. Inserts were cloned using Golden Gate cloning with BsaI (NEB) into a pET22 plasmid. Plasmids were transformed into E. coli DH5 alpha and sequenced confirmed by Sanger sequencing through Functional Biosciences.

For expression of the assembled lysins, plasmids were transformed into E. coli BL21 (DE3). Starter cultures were made by inoculating from glycerol stocks into 5mL of LB medium containing 100μg/mL Carbenicillin and cultured overnight at 37°C and 250 rpm. 500μL of starter cultures were used to inoculate 50mL of Terrific broth in 250mL baffled flasks. Cultures were grown at 37°C at 250rpm to an OD600 of ~0.6. Protein expression was induced with 50μM IPTG, and cultures were transferred into a pre-chilled shaker at 20°C and grown for 18 hours at 250rpm. Cultures were then harvested by centrifugation in a tabletop centrifuge at 3000xG for 10 minutes at 4°C.

### Growth inhibition assays

All experiments were preceded by thoroughly sterilizing the workspace and material with spore-klenz and subsequently with 70% ethanol to avoid contamination of bacteria and bacterial spores. For each experiment, precultures of each species were prepared by thawing a single-use glycerol stock and combining the inoculation volume and media to a total volume of 10 mL until the stationary phase was reached at 37°C. Incubation times are also listed in **Supplementary Table 2**. The OD600 values of the precultures were measured (Tecan F200 Plate Reader, 200 μL in 96-Well Microplate) and used to normalize each culture to an OD600 of 0.0066 by spinning down at 3200 x g for 10 min and resuspending with DM38. E coli lysate containing expressed lysins was prepared by pelleting the expression culture at 3,000xg for 5 minutes, discarding the supernatant, and resuspending in 1x PBS (137 mM NaCl, 2.7 mM KCl, 10 mM Na_2_HPO_4_, 1.8 mM KH_2_PO_4_) at 1/10 of the culture volume. Resuspended cultures were then lysed by sonication (5 seconds on 15 seconds off for a total on time of 1 minute. Sonicated samples were centrifuged at 4,000xg at 4°C for 30 minutes and the soluble fraction was transferred into conical tubes and stored on ice for temporary storage. Samples and controls were prepared by adding concentrated lysate to normalized pre-culture in 96-well plates to a final volume of 200μL. Concentrated empty-plasmid lysate was used as controls for each strain tested and each condition was done in triplicate. Plates were then sealed using semi-permeable membranes (Breathe-Easy seal Sigma-Aldrich) and placed on a microplate reader (Tecan F200 Plate Reader). OD600 was then monitored for 24-48 hours, for fast and slow growing strains respectively and shown in **Supplementary Table 2**, at 37°C in a plate reader. With the data obtained we calculated growth inhibition for each sample following the equation:

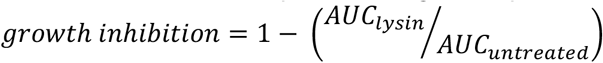

Where *AUC*_*lysin*_ is the area under the growth curve for the lysin treated sample and *AUC*_*untreated*_ is the under the growth curve for the untreated sample. Ridge regression was performed on the growth inhibition results using RidgeCV method with default parameters from the scikit-learn library in Python3 to estimate the domain contributions to growth inhibition.

For single strain titrations, each purified protein was first diluted to 2μM in 100mM Tris/HCl, 150mM NaCl pH 8.0 buffer. Then a 2-fold series dilution was performed such that final assay concentrations were 200nM, 100nM, 50nM, 25nM, 12.5nM, 6.25nM and 3.125nM, including a 0nM buffer negative control. All samples and control conditions were done in triplicate.

### Protein purification

Cells were grown at 37°C, 250RPM, to OD600 0.55-0.65 and induced with 100μM IPTG. Induced cultures were then grown at 30°C, 250RPM, for 14-16 hours. After induction growth, cells were harvested by centrifugation at 3000xG, and the supernatant was discarded. Cell pellets were resuspended in 100mM Tris/HCl, 150mM NaCl pH 8.0, and lysed by sonication (5 seconds on, 15 seconds off, total sonication time 1 minute). Cell lysates were clarified by centrifugation at 4,000xG for 1 hour. Strep-tagged clarified lysates were purified by passing through a gravity column with Strep-tactin Sepharose resin (IBA), and proteins were eluted using 100mM Tris/HCl, 150mM NaCl, 2.5 mM desthiobiotin, and analyzed by SDS-PAGE. Buffer exchange was performed by dialyzing overnight in 100mM Tris/HCl and 150mM NaCl and was concentrated using 30kDa MW (Vivaspin) protein concentrator columns 30kDa MW. Stocks were aliquoted, flash-frozen in liquid nitrogen, and stored at −80°C. Protein concentrations were determined using a Bradford assay from fresh, concentrated stock.

### Strain maintenance and anaerobic culture conditions

All anaerobic culturing was carried out in an anaerobic chamber (Coy Labs) with an atmosphere of 2.5 ± 0.5% H2, 15 ± 1% CO2, and balance N2. All prepared media, stock solutions, and materials were placed in the anaerobic chamber and left overnight to equilibrate. The strains used in this work were obtained from the sources listed in **Supplementary Table 2**, and permanent stocks of each were stored in 25% glycerol at −80 °C. Batches of single-use glycerol stocks were produced for each strain by first growing a culture from the permanent stock in anaerobic basal broth (ABB) media (HiMedia or Oxoid) to stationary phase, mixing the culture in an equal volume of 50% glycerol and aliquoting 400 μL into Matrix Tubes (ThermoFisher) for storage at −80 °C. Quality control for each batch of single-use glycerol stocks included (1) plating a sample of the aliquoted mixture onto LB media (Sigma-Aldrich) for incubation at 37 °C in ambient air to detect aerobic contaminants and (2) Illumina sequencing of 16S rDNA isolated from pellets of the aliquoted mixture to verify the identity of the organism.

### Community culturing and lysin titration

All experiments were preceded by thoroughly sterilizing the workspace and material with spore-klenz and subsequently with 70% ethanol to avoid contamination of bacteria and bacterial spores. We used defined media with components listed in **Supplementary Table 3**. For each experiment, precultures of each species were prepared by thawing a single-use glycerol stock and combining the inoculation volume and media to a total volume of 10 mL until the stationary phase was reached at 37°C. Incubation times are also listed in **Supplementary Table 2**. The OD600 values of the precultures were measured (Tecan F200 Plate Reader, 200 μL in 96-Well Microplate) and used to normalize each culture to an OD600 of 0.0066 by spinning down at 3200 x g for 10 min and resuspending with DM38.

The tested communities were combined in a 96 deep-well plate by mixing equal volumes of each species’ diluted preculture. Lysins were added to the communities to final concentrations of 0, 0.01, 0.1, and 1 uM. The 96 deep well plate was covered with a semi-permeable membrane and incubated at 37 °C for 48 h. After the incubation, 20 uL of each sample was diluted in 96-Well Microplate with 180 uL PBS buffer for OD600 measurement. 200 μL of each sample was transferred to a new 96DW plate and pelleted by centrifugation at 3200×g for 10 min. 180 uL of each supernatant was removed from each sample and stored at −20 °C for subsequent metabolite quantification by high performance liquid chromatography (HPLC). The remaining cell pellets were stored at −80 °C for subsequent genomic DNA extraction for 16S sequencing.

### HPLC quantification of organic acids

Supernatant samples were thawed at room temperature and added 2 μL of H2SO4. The samples were then centrifuged at 2400 × g for 10 min, and then 150 μL of each sample was filtered through a 0.2 μm filter using a vacuum manifold before transferring 70 μL of each sample to an HPLC vial. HPLC analysis was performed using either a ThermoFisher (Waltham, MA) Ultimate 3000 UHPLC system equipped with a UV detector (210 nm) or a Shimadzu HPLC system equipped with a SPD-20AV UV detector (210 nm). Compounds were separated on a 250 × 4.6 mm Rezex© ROA-Organic acid LC column (Phenomenex Torrance, CA) run with a flow rate of 0.2 mL min−1 and a column temperature of 50 °C. The samples were held at 4 °C before injection. Separation was isocratic with a mobile phase of HPLC grade water acidified with 0.015 N H2SO4 (415 μL L−1). At least two standard sets were run along with each sample set. Standards were 100, 20, and 4 mM concentrations of butyrate, succinate, lactate, and acetate. For most runs, the injection volume for both sample and standard was 25 μL. The resultant data was analyzed using the Thermofisher Chromeleon 7 software package or the Shimadzu LabSolutions software package.

### Genomic DNA extraction, DNA library preparation, sequencing, primer design, and data analysis

DNA extraction, library preparation, and sequencing were performed according to methods described in Hromada 202115 and Clark 2021 16. After experiments, cell pellets from 200 uL of culture were stored at −80C. Genomic DNA was extracted using a version of the DNeasy protocol (Qiagen) adapted for 96-well plates. Genomic DNA was normalized to 1 ng/μL in molecular-grade water and stored at −20°C. Dual-indexed primers for multiplexed amplicon sequencing of the v3-v4 region of the 16S gene were designed as described previously and arrayed in 96-well, skirted PCR plates (check brand) using an acoustic liquid handling robot (confirm with Venturelli lab). Genomic DNA and PCR master mix were added to primer plates and amplified before sequencing on an Illumina MiSeq platform. Sequencing data were analyzed as described in Hromada 2021. In brief, basespace Sequencing Hub’s FastQ Generation demultiplexed the indices and generated FastQ files. Paired reads were merged using PEAR (Paired-End reAd mergeR) v0.9.0 (Zhang et al., 2014)67. Reads 500 were mapped to a reference database of species used in this study, using the mothur v1.40.5 and the Wang method (Wang et al., 2007; Schloss et al., 2009)68,69. Relative abundance was 502, calculated by dividing the read counts mapped to each organism by the total reads in the sample. 503 Absolute abundance was calculated by multiplying the relative abundance of an organism by the 504 OD600 of the sample. Samples were excluded from further analysis if > 1% of the reads were 505 assigned to a species not expected to be in the community (indicating contamination)

## Supplementary Information

**Supplementary Figure 1.**
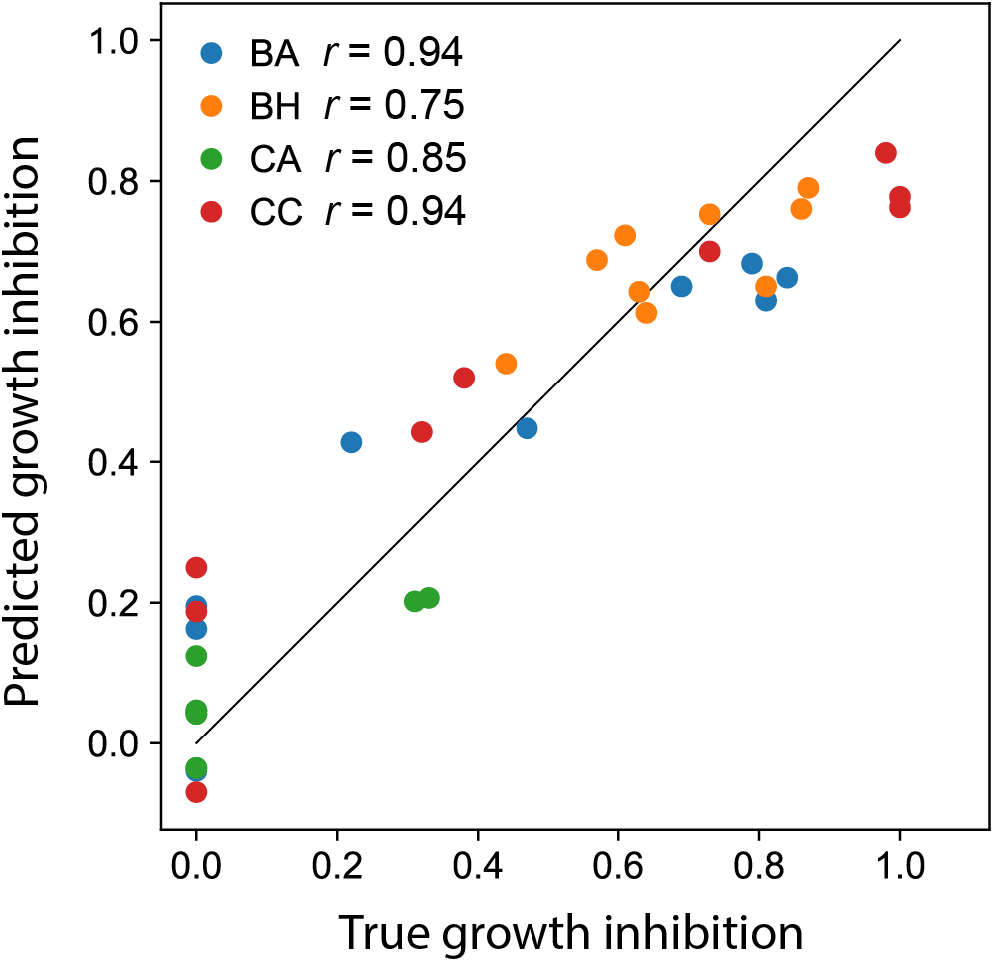
Linear regression model fit. We used ridge regression to estimate the domain contributions to growth inhibition on the four species tested. The scatter plot shows the results from leave-one-out cross validation showing a linear additive model is describes the chimeric lysin activity well.

